# Nicotine enhances intravenous self-administration of cannabinoids in adult rats

**DOI:** 10.1101/2022.10.06.510908

**Authors:** Sierra J. Stringfield, Bryson E. Sanders, Jude A. Suppo, Alan F. Sved, Mary M. Torregrossa

**Affiliations:** Department of Psychiatry, University of Pittsburgh, Pittsburgh, PA, USA; Department of Neuroscience, School of Arts and Sciences, University of Pittsburgh, Pittsburgh, PA, USA; Center for Neuroscience, University of Pittsburgh, Pittsburgh, PA, USA

**Author notes:** Corresponding author: Mary M. Torregrossa, Department of Psychiatry, University of Pittsburgh, 450 Technology Dr. Suite 223, Pittsburgh, PA 15219, United States. These authors contributed equally to this work (Alan F. Sved, Mary M. Torregrossa).

## Abstract

**Introduction:** Nicotine and cannabis are commonly used together, yet few studies have investigated the effects of concurrent administration of both drugs. Nicotine exhibits reinforcement enhancing effects by promoting the reinforcing properties of stimuli including other drugs. As many studies of this effect have used non-contingent nicotine, we implemented a dual-self-administration model where rats are given simultaneous access to two drugs and choose which to self-administer throughout a session. Here, we investigated the effect of self-administered or non-contingently delivered nicotine on cannabinoid self-administration.

**Methods:** Adult male rats were allowed to self-administer the synthetic cannabinoid WIN 55,212-2 (WIN) intravenously, with or without subcutaneous nicotine injections before each session. A separate group of animals were allowed to self-administer WIN, nicotine, or saline using a dual-catheter procedure, where each solution was infused independently and associated with a separate operant response. A third group of male and female rats were allowed to self-administer delta-9-tetrahydrocannabinol (THC) with or without pre-session injections of nicotine.

**Results:** Nicotine injections increased self-administration of WIN and THC. During dual self-administration, nicotine availability increased saline and WIN infusions but nicotine intake was not changed by WIN or saline availability. Rats preferred nicotine over saline, but preferred nicotine and WIN equally when both were available. The effect of nicotine on cannabinoid self-administration was acute and reversible when nicotine was no longer present.

**Conclusions:** These results expand our understanding of the ability of nicotine to enhance reinforcement of other drugs of abuse and suggest that co-use of nicotine and cannabinoids promotes cannabinoid use in excess of what would be taken alone.

**Implications:** This study utilizes a dual intravenous self-administration model to investigate the ability of nicotine to enhance cannabinoid intake. Our results demonstrate that the reinforcement enhancing properties of nicotine on drug use extend to include cannabinoids, but that this effect occurs specifically when nicotine is administered alongside the cannabinoid. Interestingly, cannabinoid use did not promote nicotine intake, suggesting this mechanism of reinforcement is specific to nicotine.

## INTRODUCTION

Most drug users will have experience with more than one substance throughout their lives, yet the consequences of poly-substance use are understudied. Nicotine and cannabis are two of the most widely used substances in the United States^1^, and the majority of tobacco users also have experience with cannabis^2–4^. Both nicotine dependence and cannabis use disorder can result in adverse health consequences, and co-use of these substances may result in additional negative outcomes^4–6^.

Nicotine is the primary reinforcing compound in tobacco products and it exhibits significant reinforcement enhancing effects by increasing behavioral responding for stimuli that are not associated with acquiring nicotine itself ^7–10^. Nicotine enhances responding for non-drug stimuli, and promotes intake of other drugs of abuse such as cocaine, methamphetamine, and ethanol^11–19^. Despite these effects on other commonly abused substances, there is a substantial gap in knowledge concerning the effects of nicotine on cannabinoid reinforcement.

As polydrug use can be difficult to model in animals, we sought to implement a procedure for concurrent intravenous self-administration of two separate drugs and measure the reinforcement enhancing effects of nicotine on cannabinoid self-administration. Despite the substantial use of cannabis and cannabinoid products in humans^1^, reliable cannabinoid self-administration has been difficult to achieve in rodent models. Building on renewed interest in investigating cannabinoid effects, recent studies have worked to develop protocols for intravenous self-administration that promote cannabinoid intake and result in behavioral effects that align with humans^20–25^. We chose to adapt a model for THC self-administration that results in modest amounts of cannabinoid intake^21,24,25^ to investigate the ability of nicotine to promote cannabinoid intake in excess of the amounts that the animals would administer alone.

In the present study, we investigated the ability of nicotine to influence the acquisition and maintenance of cannabinoid self-administration. We hypothesized that concurrent nicotine exposure, either through non-contingent or volitional administration, would facilitate cannabinoid self-administration and increase intake. We found that nicotine enhanced self-administration of both WIN55,212-2 (WIN) and THC but access to these substances did not impact nicotine self-administration.

## METHODS

### Animals

Male and female Sprague Dawley rats (approximately 250 g and 225 g, respectively, on arrival) were purchased from Envigo (Indianapolis, IN). Rats were individually housed in a temperature and humidity-controlled room on a 12-hour reverse light/dark cycle (lights off at 7:00 am). Access to food (LabDiet Rodent 5001; LabDiet, St. Louis, MO) and water was provided *ad libitum*. All animal procedures were conducted in accordance with the guidelines of the National Institutes of Health *Guide for the care and use of Laboratory Animals* and were approved by the University of Pittsburgh Institutional Animal Care and Use Committee.

### Drugs

Nicotine hydrogen tartrate salt (MP Pharmaceuticals, Solon, OH) was dissolved in 0.9% sterile saline. All nicotine doses are expressed as free base. WIN55,212-2 mesylate (WIN, Cayman Chemicals, Ann Arbor, MI) or Δ-9-tetrahydrocannabinol (THC, generously provided by the National Institute on Drug Abuse’s Drug Supply program) was added to 100–200 μl of Tween 80 depending on the dose of THC. Ethanol was evaporated from THC solutions using nitrogen gas^25^. Additional description of drug preparation is provided in Supplemental Methods.

### Apparatus

Self-administration sessions occurred in standard operant chambers within sound-attenuated cubicles (Med-Associates, St Albans, VT). Chambers were configured with two nose poke ports with white stimulus lights located directly above each port. A red houselight was located on the same wall. Syringe pumps were connected to an infusion swivel with tubing to connect to catheter ports. During dual self-administration experiments, two separate syringe pumps were connected to a dual channel fluid swivel to allow for access to two distinct solutions during each session.

### Surgical Procedures

Catheters for single and dual intravenous self-administration experiments were constructed in house, additional information of catheter construction is supplied in Supplemental Methods. Rats recovered from surgery for at least 5 days. Catheters were flushed daily with 0.1 ml saline containing 30 U/ml heparin and 66.67 mg/ml gentamicin. Catheter patency was confirmed periodically with methohexital (5 mg/kg) and after the final self-administration session. A total of 18 animals with catheters that lost patency were excluded from the experiment.

### Intravenous Self-Administration Sessions

Rats completed daily 1-hour self-administration sessions. Rats in Experiments 1 and 3 self-administering one drug solution were randomly assigned to active and inactive nose poke ports, and responses into the inactive port were recorded but did not have programmed consequences. Responding in the active port resulted in the intravenous delivery of drug solution and 1 s illumination of the stimulus light located directly above the port, followed by a mandatory 60 s timeout where responses into the active port were recorded but did not result in drug infusion. For dual self-administration sessions in Experiment 2, both nose poke ports were active and resulted in an infusion of a specific solution, followed by the timeout period.

#### Experiment 1: Nicotine enhancement of WIN 55,212-2 self-administration

In Experiment 1A, male rats (N=15) were allowed to self-administer 12.5 μg/kg/infusion WIN on a fixed ratio 1 (FR1) schedule of reinforcement. The schedule of reinforcement was increased to FR5 on days 8-15 and then reduced to FR3 on days 16-21. Next, these animals were given injections of saline or 0.3 mg/kg nicotine 10-minutes prior to each session for an additional 22 days of WIN self-administration sessions. This dose of nicotine was chosen for its ability to produce moderate behavioral and physiological effects^26–29^. In Experiment 1B, to investigate the effect of a continuous presence of nicotine on acquisition and prolonged WIN self-administration, a separate group of male rats were permitted to self-administer WIN for a total of 38 days. Due to a technical error, a white houselight in the operant chamber was illuminated for the first 9 days. This error was corrected, and all sessions were run under red light for the remainder of the experiment. Rats were allowed to self-administer WIN on a FR2 schedule of reinforcement for the first 12 days, the schedule was increased to FR3 on day 13 and continued for the remainder of the experiment. Rats in the NIC group (N=8) were injected with 0.3 mg/kg nicotine 10-minutes prior to each session, while rats in the SAL control group (N=6) received saline. On day 34-38 the pre-session injections were switched so that rats in the NIC group received saline, and animals in the SAL group were injected with nicotine prior to each session.

#### Experiment 2: Dual self-administration of nicotine and WIN

Male rats were implanted with dual catheters to allow for access to two separate drug solutions within the same operant self-administration session. Rats were permitted to intravenously self-administer 12.5 μg/kg/infusion WIN, 30 μg/kg/infusion nicotine, or saline in daily 1-hr sessions. Rats were randomly assigned to treatment groups where they had access to pairs of drug solutions: WIN and NIC (N=8), WIN and SAL (N=14), NIC and SAL (N=10), or SAL and SAL (N=8). For these intravenous self-administration sessions, one nose poke port was assigned to each solution and there was no inactive port. Both solutions were available throughout the 1-hour session. Responding in one port resulted in an infusion of the associated drug along with presentation of the stimulus light located directly above the corresponding port followed by a 60 s timeout. Rats self-administered for a total of 19 days on an FR1 schedule of reinforcement.

#### Experiment 3: Nicotine enhancement of THC self-administration in male and female rats

Male and female adult rats were permitted to self-administer escalating doses of THC and received pre-session injections of nicotine or saline ^25^. For the first 8 days of training, rats in the NIC group received 3 mg/kg nicotine due to mathematical error, which led to a suppression of responding (data not shown). However, once the dose was corrected responding immediately increased to levels similar to the saline group. Rats received injections of 0.3 mg/kg nicotine (N=7 female, N=6 male) or saline (N=6 female, N=6 male) prior to each session for the remainder of the experiment. Animals began with self-administration access to 3 μg/kg/infusion THC for days 1-3 of training, followed by 10 μg/kg/infusion on days 4-6, and 30 μg/kg/infusion beginning on day 7 and continuing until the end of the experiment. The drug concentration was adjusted with each change in unit dose to maintain a similar infusion volume across sessions. On days 13-17, the pre-injection solution was switched to the opposite treatment condition, where the NIC group received a pre-session injection of saline and the SAL group received nicotine. Animals were returned to the original pre-injection conditions for days 18-22. These animals then underwent one day of progressive ratio (PR) training. The PR session lasted for 4 hours, and all rats received pre-injections of nicotine or saline according to their initial group designations. These PR sessions used 30 μg/kg/infusion THC and the number of operant responses required for an infusion escalated from 3, 6, 10, 15, 20, 25, 32, 40, 50, 62, 77, 95,118, 145, 179, 219, 268, 328, 402, 492.

### Statistical Analysis

Statistical analysis was completed using GraphPad Prism version 9.1 (GraphPad Software, San Diego, CA). Drug intake across self-administration sessions was compared by repeated measures ANOVA (group × day or sex × group × day), followed by Bonferroni’s test for multiple comparisons when applicable. Differences in WIN infusions after nicotine or saline injections were compared by paired samples t-test. α = 0.05 for all analyses.

## RESULTS

### Experiment 1: Nicotine enhancement of WIN 55,212-2 self-administration

To investigate the ability of nicotine to promote cannabinoid self-administration, we first evaluated intravenous self-administration of the synthetic cannabinoid WIN on multiple schedules of reinforcement (Supplemental Figures S1A, S2A-B). Rats responded preferentially for the active port over the inactive port at all schedules with a trend toward a main effect of schedule of reinforcement (Supplemental Figure S1A). Next, the same animals were allowed to self-administer WIN on an FR3 schedule (Figure 1B). All rats received an injection of saline (s.c.) 10 minutes prior to each session on days 1-5. On days 6-15, all rats were injected with 0.3 mg/kg nicotine (s.c.) 10 min prior to the session. Rats were returned to saline injections for days 16-18, and then nicotine injections for days 19-22 of the experiment. WIN infusions received on the final day of saline (day 18), or nicotine injections (day 22) were compared (Figure 1C, Supplemental Figure 2C) and rats self-administered more WIN after receiving an injection of nicotine than after receiving an injection of saline prior to the session (t_14_=2.42, p<0.05).

**Figure 1:**
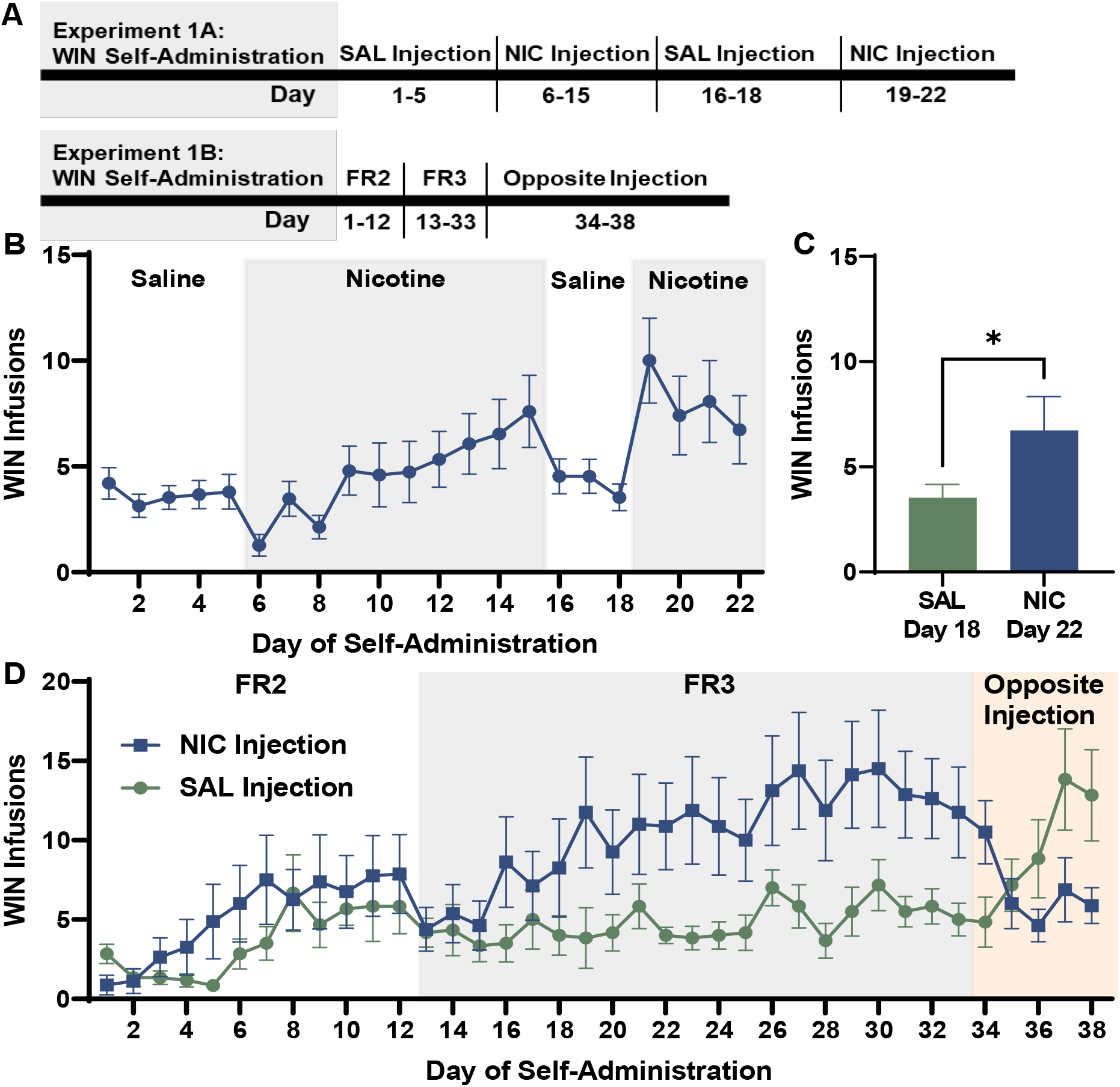
Non-contingent nicotine enhances self-administration of WIN 55,212-2. A) Timelines depicting experimental procedures. B) Mean WIN infusions from male rats (N=15) self-administering on an FR3 schedule of reinforcement after receiving pre-session injections of saline or 0.3 mg/kg nicotine. C) Mean WIN infusions received on the last day of saline injections (day 18) or nicotine injections (day 22). C) WIN self-administration in NIC (N=8) or SAL (N=6) groups receiving pre-session injections of nicotine or saline on increasing schedules of reinforcement, followed by 5 days of testing where rats received the alternate injection condition. All figures are mean (±SEM), * p<0.05.

**Figure 2:**
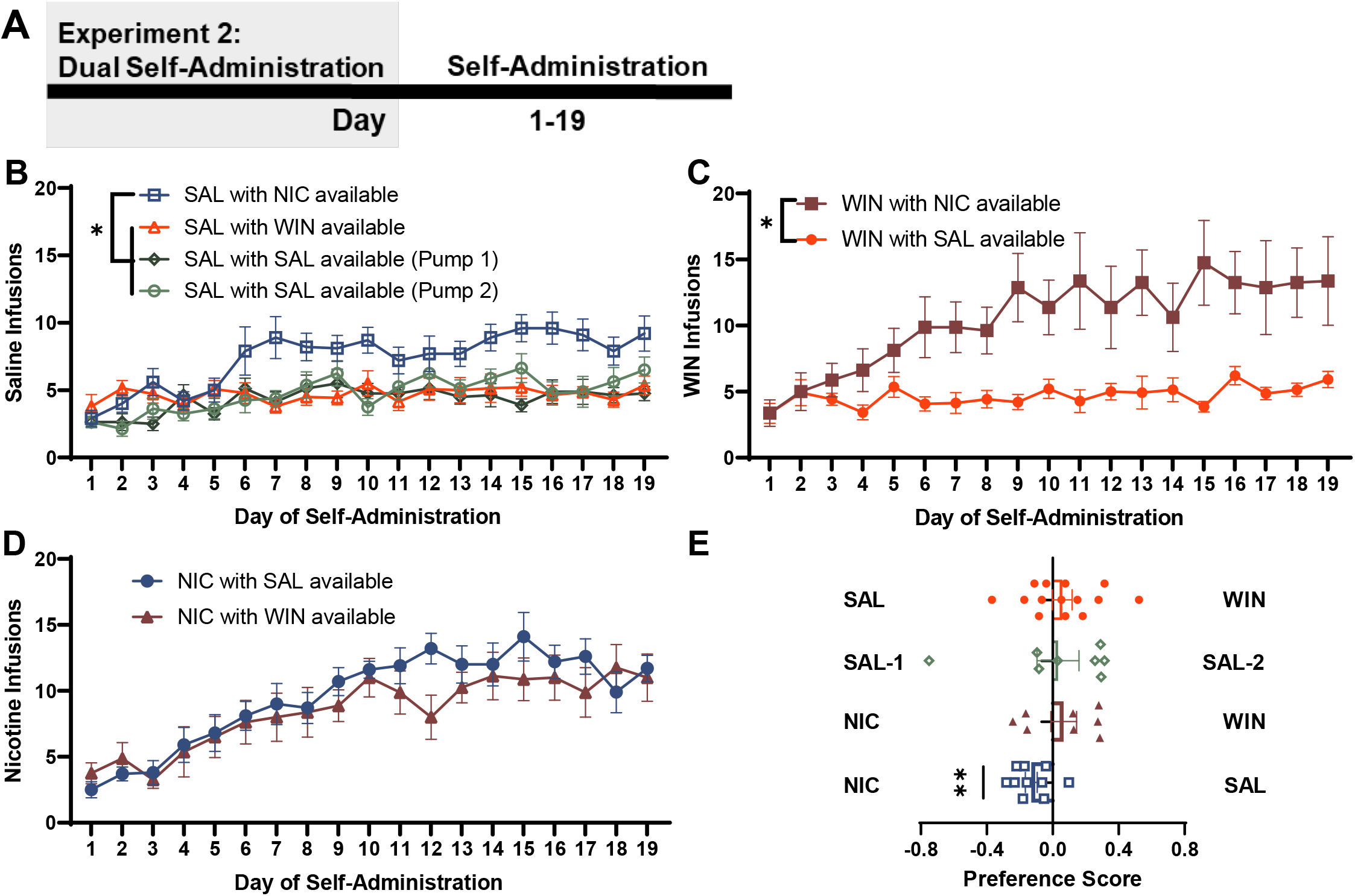
Dual self-administration of nicotine, WIN, or saline. A) Timeline of experimental procedures. B) Mean infusions per day in male rats self-administering saline (SAL) with concurrent availability of nicotine (NIC, N=10), WIN (N=14) or additional saline (N=8). C) Mean infusions of rats self-administering WIN with access to NIC (N=8) or SAL. D) Infusions received by rats self-administering NIC with access to SAL or WIN. E) Preference scores comparing the average number of infusions of each solution received over the last 3 days of self-administration. Infusions are presented as mean ± SEM, * p<0.05, ** p<0.01.

In a separate cohort of animals that consistently received nicotine (NIC) or saline (SAL) prior to every session, we investigated the ability of nicotine exposure to promote WIN self-administration when present during acquisition and if the enhancement in WIN intake would persist without nicotine exposure (Figure 1D). Rats began self-administration on an FR2 schedule of reinforcement where responding for WIN increased over time as rats acquired the behavior (main effect of day, F _(11, 132)_ = 5.95, p<0.01) but there was no main effect of group or group × day interaction (p>0.1 for all analyses). When the schedule was increased to an FR3, a day × group interaction emerged (F _(20, 240)_ = 2.05, p<0.01) along with a main effect of day (F _(20, 240)_ = 4.63, p<0.01) with rats in the NIC group receiving more WIN infusions than those in the SAL group. When the pre-session injections administered to these animals were switched, a day × group interaction emerged (F _(4, 48)_ = 9.37, p<0.001) along with a main effect of day (F _(4, 48)_ = 3.72 p<0.05). Rats in the former SAL group increased their WIN intake while rats in the former NIC group significantly reduced their WIN intake, indicating that the enhancing effects of nicotine did not persist in its absence.

### Experiment 2: Dual self-administration of WIN 55,212-2 and nicotine

We next questioned if volitional control over nicotine intake would alter WIN self-administration (Figure 2). We utilized a dual self-administration model that allowed rats to choose between two solutions available throughout the session. When saline intake was compared across rats that also had access to either nicotine, WIN, or a second saline port (Figure 2B), a day × group interaction emerged (F _(54, 648)_ = 1.77, p<0.001) as well as a main effect of day (F _(18, 648)_ = 6.84, p<0.001) and a main effect of group (F _(3, 36)_ = 9.95, p<0.001). Specifically, saline infusions were increased in animals with simultaneous nicotine availability relative to the other conditions. WIN self-administration was significantly increased when nicotine was concurrently available relative to saline (Figure 2C) as evident by a day × group interaction (F _(18, 360)_ = 5.15, p<0.001), main effect of day (F _(18, 360)_ = 7.43 p<0.01), and main effect of group (F _(1, 20)_ = 13.11, p<0.01). While concurrent availability of nicotine enhanced self-administration of both WIN and saline, access to WIN or saline did not influence nicotine self-administration (Figure 2D). Rats increased nicotine self-administration over time (main effect of day, F _(18,288)_ = 14.79, p<0.001) but no main effect of group or day × group interaction emerged (p>0.1 for all analyses). Thus, nicotine enhances responding for concurrently available WIN or saline, but these solutions do not reciprocally enhance nicotine intake.

Interestingly, animals self-administering WIN or saline with access to nicotine earned comparable numbers of WIN or saline infusions (t_16_=1.53, p>0.1). To further explore the impact of nicotine on WIN and saline self-administration, we compared individual preference scores across the last three self-administration sessions (Figure 2E). Although nicotine enhanced saline intake, rats still exhibited a significant preference for nicotine during dual self-administration sessions (t_9_=3.62, p<0.01). This preference for nicotine was not present in rats self-administering nicotine and WIN (t_7_=0.85, p>0.1). Control rats did not exhibit a preference for saline delivered on either side (t_7_=0.25, p>0.1), and rats exhibited a similar preference for saline or WIN when both were available (t_13_=0.96, p>0.1). This effect was also present when response ratios were computed to compare entries into each drug solution port (Supplemental Figure S4), where a significant difference in response ratio was only present in rats that self-administered saline and nicotine (t_18_=5.04 p<0.001). Thus, while nicotine enhanced the amount of both saline and WIN the animals chose to take, the solutions were not equal in their value as reinforcers in the presence of nicotine.

### Experiment 3: Nicotine enhancement of THC self-administration in male and female rats

As WIN is a synthetic cannabinoid not used for human consumption, we next investigated if nicotine would produce similar reinforcement enhancing effects on self-administration of THC, the primary psychoactive component of cannabis. Male and female rats were allowed to self-administer THC after receiving pre-session injections of nicotine (NIC group) or saline (SAL group, Figure 3). No significant difference in THC intake emerged between male and female rats for the number of active port entries (sex × day × group interaction F _(21, 441)_ = 0.66, p>0.1), inactive port entries (sex × day × group interaction F _(21, 441)_ = 1.07, p>0.1) or infusions (Figure 3B, sex × day × group interaction, F _(21, 441)_ = 0.95 p>0.1). Thus, drug infusions were collapsed across sex to evaluate the effect of passive nicotine injection on THC self-administration. Rats that received pre-injections of nicotine showed increased THC self-administration compared to those that received saline injections as evident by a main effect of group, (F _(1, 23)_ = 11.19, p<0.01) a main effect of day (F _(11,253)_ = 5.98, p<0.001), and a day × group interaction (F _(11, 253)_ = 4.45, p<0.001). While nicotine pre-injections promoted responding for THC, there was no difference in response ratio between groups on days 10-12 of self-administration (Supplemental Figure S1C). When injection conditions were switched THC intake changed to reflect this change resulting in a main effect of group (F _(1, 23)_ = 5.96, p<0.05) and a group by day interaction (F _(4, 92)_ = 9.63, p<0.01). When injection conditions returned to the original groups, THC intake recovered with animals receiving nicotine injections increasing their intake, and animals receiving saline injections reducing their intake as evident by a main effect of group (F _(23, 92)_ = 5.32 p<0.01) and a trend toward a day by group interaction (F _(4, 92)_ = 2.35 p=0.06). Thus, like self-administration of the synthetic cannabinoid WIN, nicotine exhibits acute reinforcement enhancing effects on THC self-administration in both male and female rats.

**Figure 3:**
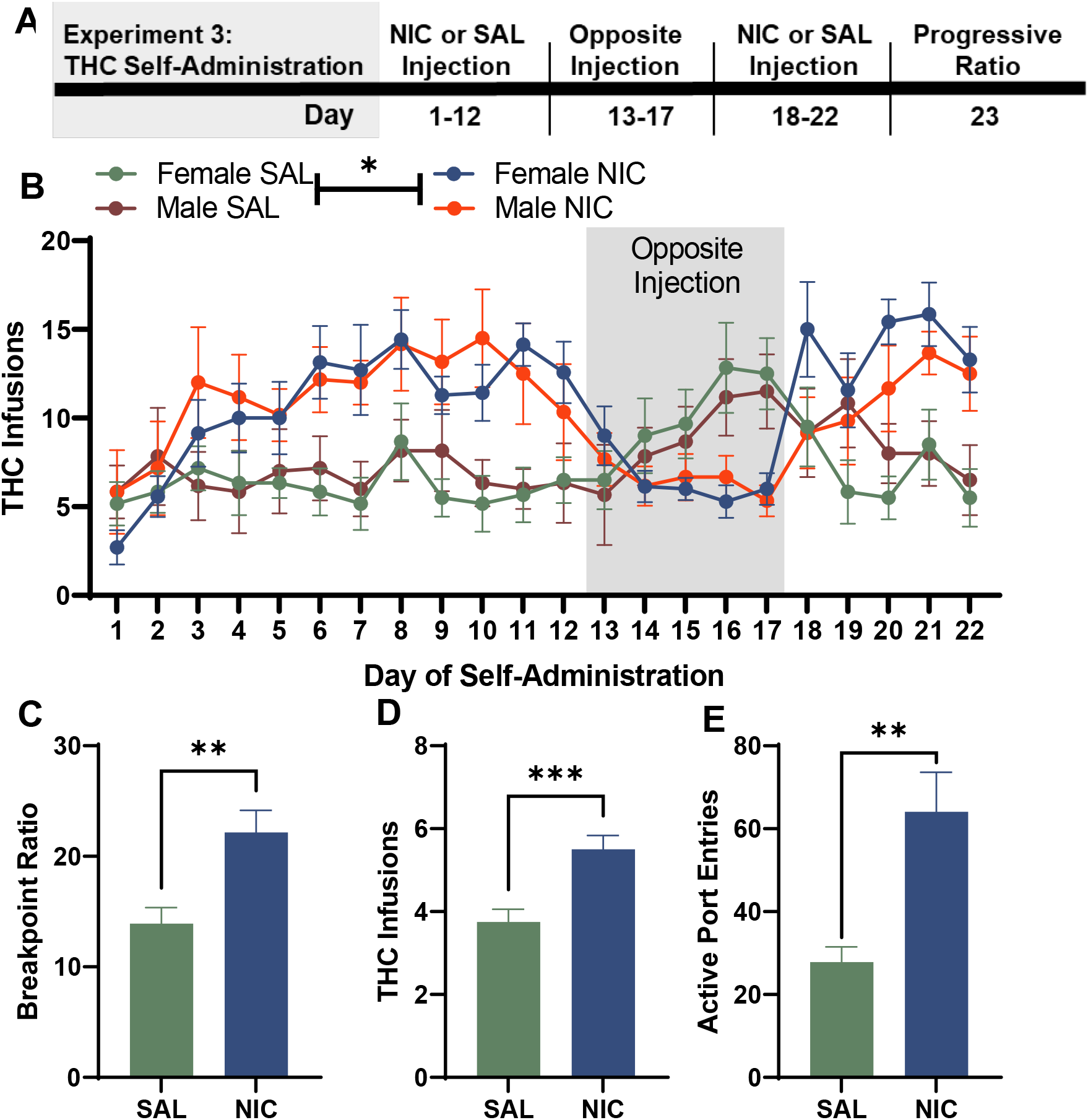
Nicotine enhancement of THC self-administration in male and female rats. A) Timeline of experimental procedures. B) Mean infusions in rats receiving pre-session injections of nicotine (NIC N=7 female, N=6 male) or saline (SAL, N=6 female, N=6 male) prior to intravenous THC self-administration. Rats received the alternate injection condition over a 5-day period, before returning to the original injection conditions. C) Average final ratio completed over 4 days of progressive ratio testing for male and female rats self-administering THC after pre-session injections of nicotine or saline. D) Mean infusions received for each group on each day of progressive ratio training. E) Average number of active port entries across each day of progressive ratio training. * p<0.05, ** p<0.01, *** p<0.001.

Next, the same cohort of rats completed one day of progressive ratio testing (Figure 3C-E). As no significant difference in responding emerged between males and females on any measure tested, results were collapsed across sex to compare the effect of nicotine pre-injection on willingness to respond for THC. Animals in the NIC group achieved a higher breakpoint or final ratio of responses to receive an infusion of THC (Figure 3C; t_22_=3.35, p<0.01). They also received more infusions (Figure 3D; t_22_=3.85, p<0.001) and completed more active port entries than animals in the SAL group (Figure 3E; t22=3.55, p<0.01). These results indicate that nicotine not only promoted THC self-administration but promoted drug-intake under increasingly demanding response requirements.

## DISCUSSION

In the present study, we demonstrate that the reinforcement enhancing effects of nicotine extend to self-administration of the cannabinoid WIN in male rats, and THC in males and females. Using a dual self-administration procedure in male rats, we found that nicotine promoted intake of both WIN and saline, but access to these solutions did not impact nicotine intake. Rats preferred to receive more nicotine infusions when saline was the alternative but chose to receive comparable infusions of nicotine and WIN when both were available. The enhancing effect of nicotine was only present while nicotine was onboard, as drug intake fell to control levels when the animals’ received injections of saline or self-administered saline. The ability of nicotine to serve as a primary reinforcer, while also promoting seeking of reinforcers unassociated with nicotine itself has been well established ^8,10,30,31^, and these effects apply to non-drug stimuli and drugs of abuse ^12,18^. The present results extend this effect to cannabinoids, further increasing understanding of the impact of nicotine on drug use.

In the dual self-administration experiment, male rats self-administered modest amounts of WIN on par with the saline control group. Previous studies have indicated a need for food restriction, food training, prior vapor exposure, or escalation of doses to establish intravenous cannabinoid intake, and we did not employ any of these methods to promote self-administration in this study^20,21,25,32^. Interestingly, this experience aligns with nicotine self-administration where despite consistent human use and dependency, intravenous self-administration was notoriously difficult to achieve in the absence of a visual stimulus^8,33^. Recent studies focused on developing rodent models of intravenous cannabinoid intake have encountered similar problems in achieving reliable intake, particularly when compared to expectations for stimulant or opioid self-administration. Self-administration of cannabinoid extracts using a vapor inhalation model, as well as models that promote oral intake of cannabinoids through drinking or food have been successfully applied to rodents^34–37^. Despite these indications that cannabinoids can be rewarding enough to produce voluntary intake, intravenous self-administration of cannabinoids has been tenuous. Substances that promote robust intravenous self-administration in rodents also promote high levels of dependency in humans compared to cannabinoids, and the low rate of self-administration for cannabinoids aligns with expectations based on human patterns of use. Intravenous models can succeed at promoting cannabinoid intake that produce blood plasma concentrations or behavioral effects that align with expectations of human use^20^.

The enhancing effects observed during dual self-administration experiments were not bi-directional as self-administration of nicotine increased WIN intake, but access to WIN did not promote nicotine self-administration. The lack of this reciprocal effect diverges from previous work where prior exposure to THC promoted nicotine self-administration ^38^. Here, we tested concurrent WIN and nicotine self-administration, while the prior study used a higher dose of THC that was given separate from nicotine self-administration. These procedural differences may account for the difference in nicotine self-administration, as the higher dose of THC may have resulted in alterations to endocannabinoid or neurotransmitter signaling that contributed to enhanced drug-seeking over time.

We also found that nicotine enhanced saline self-administration, but interestingly, animals that increased their saline intake still preferred to self-administer nicotine when both solutions were available. This differs from animals self-administering WIN and nicotine, where the presence of nicotine enhanced WIN intake and led to no significant preference for WIN versus nicotine. In animals self-administering both WIN and saline, we found no difference in preference for either solution, reflective of the low rates of reinforced responding found in previous research with adult intravenous cannabinoid self-administration ^20,21^. While WIN alone was slightly reinforcing, the presence of nicotine promoted WIN intake and the ability of WIN to compete with nicotine as a reinforcer, an effect that did not occur with saline.

This study focuses on the effect of concurrent administration of nicotine and cannabinoids in adult animals. The fact that these experiments were primarily performed in males is a limitation of the study, although we found no difference between male and female animals during THC self-administration sessions. Females have previously been reported to self-administer more WIN than males and several studies indicate that females will self-administer elevated amounts of other drugs of abuse^39^. However, sex differences in drug self-administration are dose-dependent and prior studies have not used the same THC self-administration procedures used here. Thus, the possibility remains that sex-differences in cannabinoid self-administration could emerge but were not observed in this study. Replicating the dual self-administration experiment in females would be of particular interest, as gonadal hormone activity may be relevant for the rewarding properties of cannabinoids and these effects may interact with the reinforcement enhancing properties of nicotine^40^. While we demonstrate that the enhancing effects of nicotine on drug intake are acute and limited to instances when nicotine is present during behavioral sessions, long-term effects on neurobiological processes or behaviors may still emerge. Future studies can investigate whether nicotine exposure is sufficient to produce relapse-like behavior and reinstatement of cannabinoid seeking, similar to previous research with cocaine ^11^. Additional studies can also examine the mechanism by which nicotine co-use facilitates increased self-administration, and if it is a similar effect of nicotine across drugs of abuse. Nicotine enhancement of alcohol self-administration relies on the release of dopamine in the striatum ^41,42^ and co-exposure of nicotine and other abused substances results in alterations to striatal structure and function ^43^, suggesting a circuit of interest that may contribute to the effect of nicotine on cannabinoid self-administration.

With this study, we demonstrate that nicotine exerts reinforcement enhancing effects on cannabinoid self-administration. This result is relevant to understanding adverse outcomes associated with substance co-use or abuse. The acute enhancement of cannabinoid intake in the presence of nicotine is important for the potential health effects of increased tobacco or cannabis administration, and these effects of concurrent use can continue to be studied using preclinical models.

## Funding

This work was supported by the National Institutes of Health (grant number P50DA046346 to [AS and MT], R21DA052663 to [MT and AS], R01DA042029 to [MT], F32DA047029 to [SS], and R21DA046336 to [AS]).

## Declaration of Interests

The authors declare no conflicts of interest.

## Acknowledgements

The authors would like to thank Jennifer Zeak for outstanding technical assistance.

## Data Availability Statement

The data underlying this article will be shared on reasonable request to the corresponding author.

## Notes

### Competing Interest Statement

The authors have declared no competing interest.

