## Supplemental Materials for "Nicotine enhances intravenous self-administration of cannabinoids in adult rats"

### Supplemental Methods

#### Drug Preparation and Self-Administration

THC and WIN solutions were suspended in 0.9% sterile saline and prepared fresh weekly. Nicotine solutions were prepared fresh every 2-4 weeks and stored at 4 °C. Solutions were vortexed and drug syringes were prepared daily and sterile filtered immediately before each session. To ensure each animal received the correct dose of drug each day, the infusion volume for drug or saline control solutions was adjusted based on each animals' body weight taken immediately before each session.

#### Catheter Construction

Catheters were constructed using silastic tubing attached to a threaded, plastic pedestal bolt fitted with stainless steel tubing through the middle and bonded to surgical mesh. Double-lumen catheters were constructed from two pedestals spaced 1 cm apart, attached to separate lengths of silastic tubing. Rats were anesthetized using isoflurane and indwelling catheters were implanted into the right jugular vein for single solution self-administration (Experiments 1 and 3) and into the right and left jugular veins for dual solution self-administration (Experiment 2).

#### Statistical analysis

Supplemental analysis was completed to compare changes in drug intake or responses across schedules of reinforcement and drug exposure conditions, where mean responses over the final 3 days of each manipulation were analyzed by unpaired t-test, paired t-test, or one-way repeated measures ANOVA followed by Tukey's test for multiple comparisons when applicable. Preference scores for comparing active and inactive port entries or solution preference during dual self-administration were computed using the formula $\frac{Response_1 - Response_2}{Total\ Responses}$ . Analysis of infusions completed within the final session of self-administration for each experiment was completed by collapsing the number of infusions into 5 or 15-minute bins for each animal. Within-session analysis was analyzed by 2- or 3-way repeated measures ANOVA (group × time or sex × group × time).

**Supplemental Results and Figures**

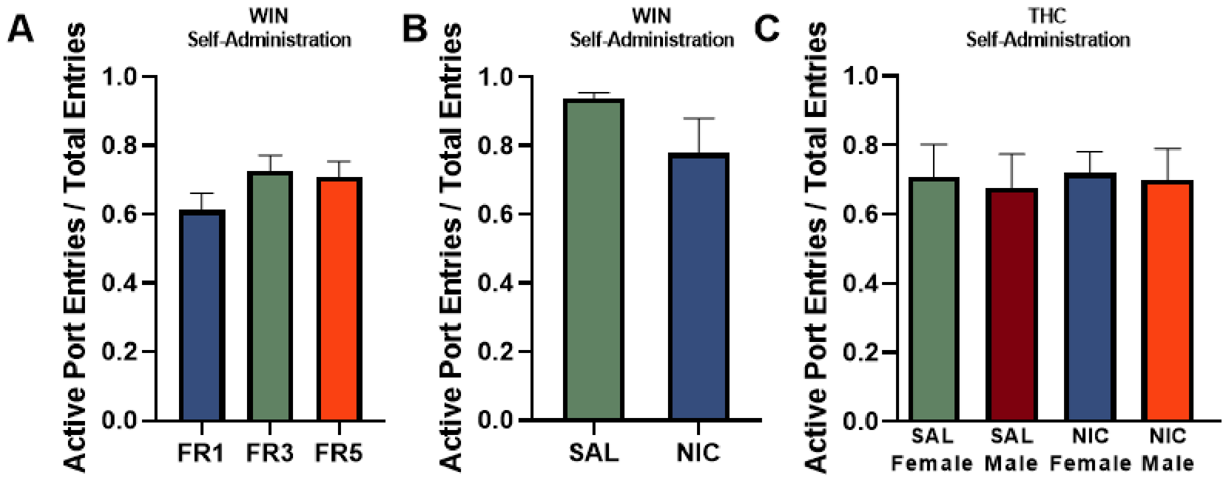


**Supplemental Figure S1**

Response ratios were computed using the formula $\frac{Responses}{Total\ Responses}$ to evaluate responses into the active port (Experiments 1 and 3) collapsed over the last 3 days of self-administration for each experiment. A) Response ratios from Experiment 1A were calculated to compare responses into the active port during each schedule of reinforcement, there was a trend toward a main effect of schedule of reinforcement, ($F_{(2, 28)} = 3.31$, p=0.051). B) When animals self-administered WIN with consistent pre-session injections of nicotine (NIC) or saline (SAL) during Experiment 1B, no differences in response ratios emerged between groups ($t_{13}$=1.26, p>0.1). C) Response ratios calculated for male and female rats self-administering THC with pre-injections of SAL or NIC, no difference between groups ($F_{(3,21)} = 0.04$, p>0.1).

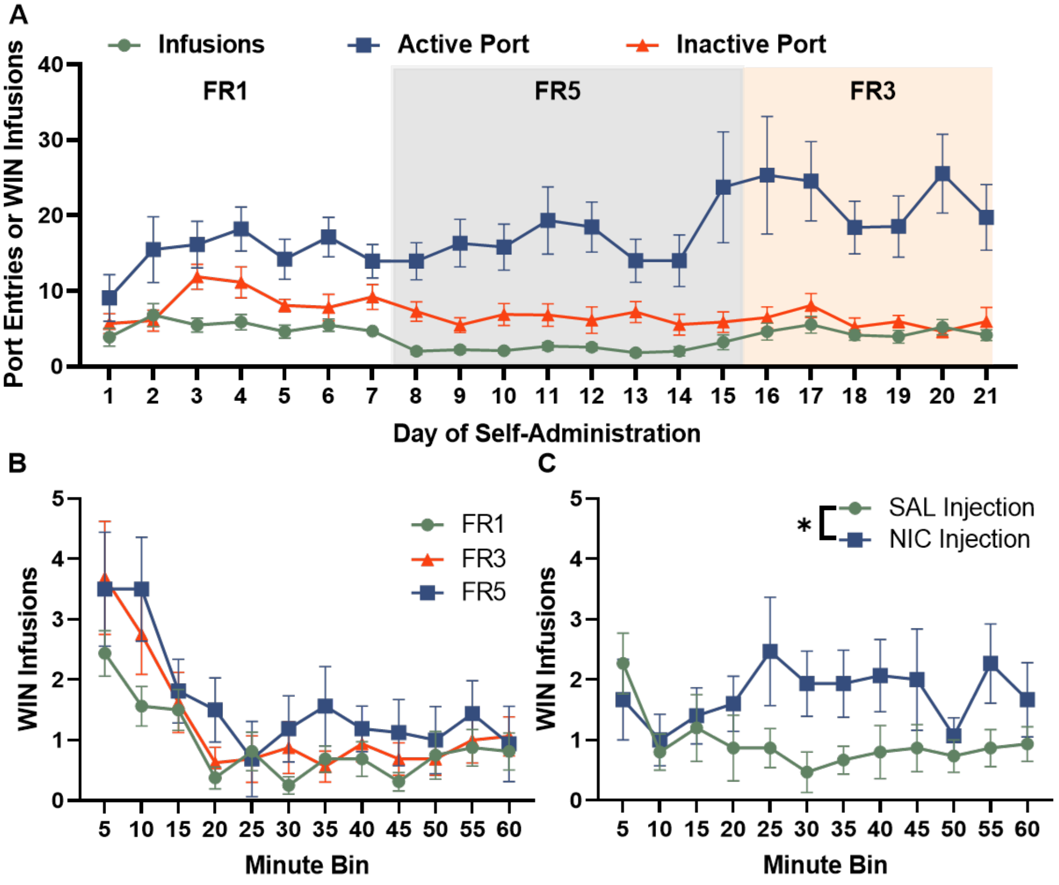


**Supplemental Figure S2**

A) Male rats self-administered WIN on FR1, FR5, and FR3 schedules of reinforcement and the number of infusions received was compared by 1-way repeated measures ANOVA. Rats received more WIN infusions on the FR1 or FR3 schedules compared to the FR5 ($F_{(2,18)} = 35.18$, $p<0.001$). B) The number of WIN infusions obtained within the final 1-hour self-administration session of each schedule of reinforcement was divided into 5-minute bins. A 2-way repeated measures ANOVA comparing responding over time at each schedule revealed a main effect of time ($F_{(11, 165)} = 11.82$, $p<0.001$), a trend toward a main effect of schedule ($F_{(2, 30)} = 3.28$, $p=0.072$), and no time × schedule interaction ($F_{(22, 330)} = 0.66$, $p>0.1$). C) Next, when the same animals received pre-session injections of saline (SAL) or nicotine (NIC) while continuing to self-administer WIN, infusions were compared over the final session of WIN self-administration with saline injections and the final session of WIN self-administration with nicotine injections. Analysis by 2-way ANOVA yielded a main effect of injection condition ($F_{(1, 14)} = 5.74$, $p<0.05$), but no main effect of time ($F_{(11, 154)} = 0.971$, $p>0.1$) nor time × injection condition interaction ($F_{(11, 154)} = 1.34$, $p>0.1$).

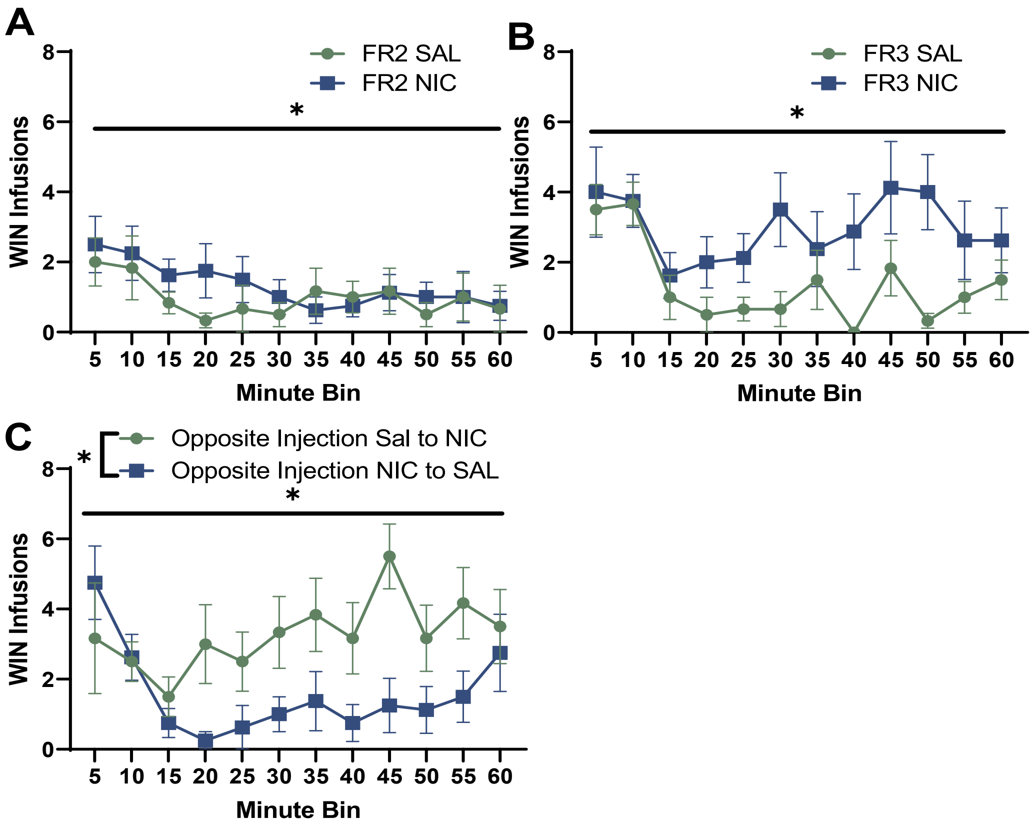


**Supplemental Figure S3**

A separate group of animals (Experiment 1B) was consistently injected with nicotine or saline prior to each WIN self-administration session. A) Comparison of within session WIN infusions between rats injected with nicotine and saline on the last day of the FR2 schedule of reinforcement (day 12) produced a main effect of time ($F_{(11, 132)} = 2.17$, $p<0.05$), but no main effect of injection condition ($F_{(1, 12)} = 0.41$, $p>0.1$) nor time × injection condition interaction ($F_{(11, 132)} = 0.62$, $p>0.1$). B) The schedule of reinforcement increased to a FR3, and within-session infusions were compared on the final day of WIN self-administration on this schedule. There was a main effect of time ($F_{(11, 132)} = 3.23$, $p<0.05$), a trend toward a main effect of injection condition ($F_{(1, 12)} = 3.62$, $p=0.081$), and no injection × time interaction ($F_{(11, 132)} = 1.2,6$ $p>0.05$). C) Injection conditions were then switched, and the time course of WIN infusions was compared on the final day of self-administration (day 38). A main effect of time ($F_{(11, 132)} = 2.56$, $p<0.05$), a main effect of injection condition ($F_{(1, 12)} = 6.17$, $p<0.05$), and a time × injection condition interaction (F (11, 132) = 2.26, $p<0.05$) emerged, where animals given nicotine injections received more WIN infusions across the session compared to animals that received saline injections that day.

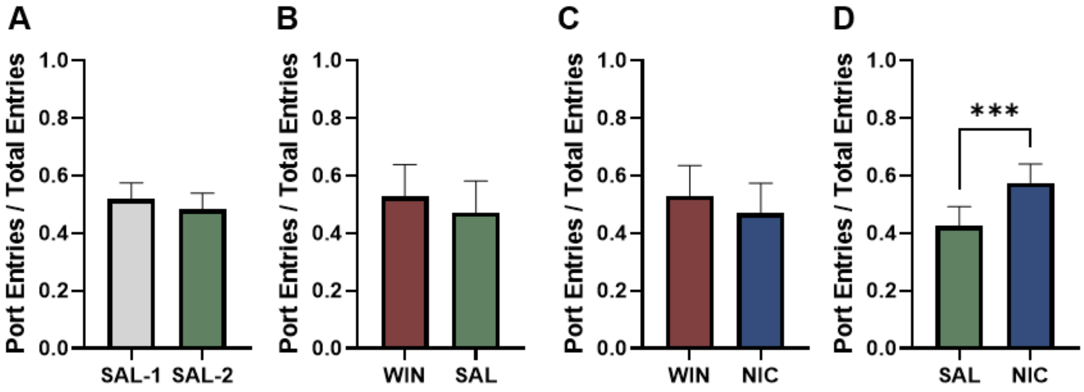


**Supplemental Figure S4**

Response ratios were computed using the formula $\frac{Responses}{Total\ Responses}$ to evaluate responses for each drug solution of interest during dual self-administration experiments (Experiment 2). A) In animals that self-administered saline (SAL) in either port, there was no effect of solution ($t_{16}$=0.45, p>0.5). B) In animals that self-administered WIN and SAL, there was no effect of solution ($t_{26}$=1.36, p>0.1). C) In animals that self-administered WIN and NIC, there was no effect of drug solution ($t_{16}$=1.25, p>0.5). D) In animals that self-administered saline and nicotine, there was an effect of solution, ($t_{18}$=5.04, p<0.05) with animals in performing more entries to receive nicotine than saline.

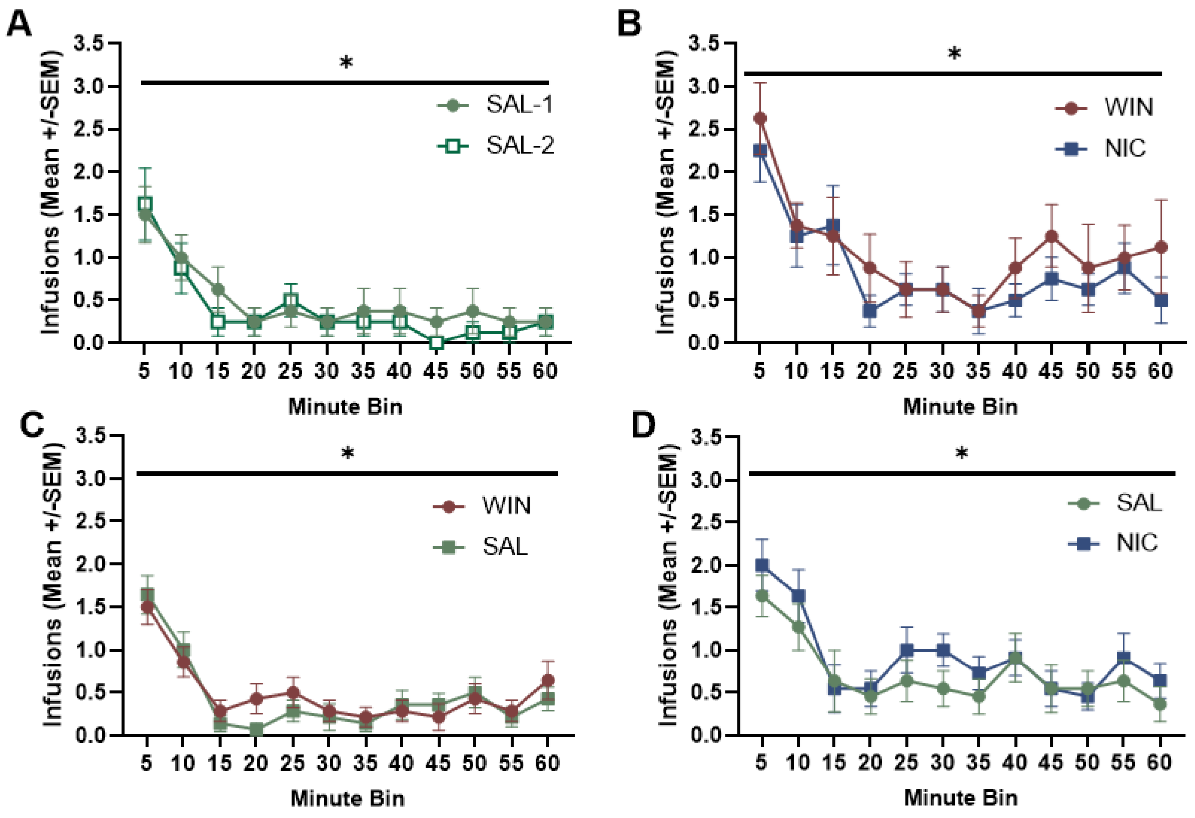


**Supplemental Figure S5**

Within-session infusions received during dual self-administration experiments. Average infusions are binned into 5-minute intervals over a 60 minute session. A) Mean infusions across the session of rats self-administering saline on both pumps, main effect of time ($F_{(11, 77)} = 6.98$, $p<0.01$). B) Mean infusions across the session of rats self-administering WIN and NIC on either pump, main effect of time ($F_{(11, 77)} = 7.570$, $p<0.01$). C) Mean infusions across the session in rats self-administering WIN and SAL on either side, main effect of time ($F_{(11, 143)} = 12.62$, $p<0.01$). D) Mean infusions across the session of rats self-administering SAL and NIC on either pump, main effect of time ($F_{(11,110)} = 4.55$, $p<0.01$), trend toward a main effect of drug solution ($F_{(1, 10)} = 4.112$, $p=0.07$). *$p<0.05$, main effect of time.

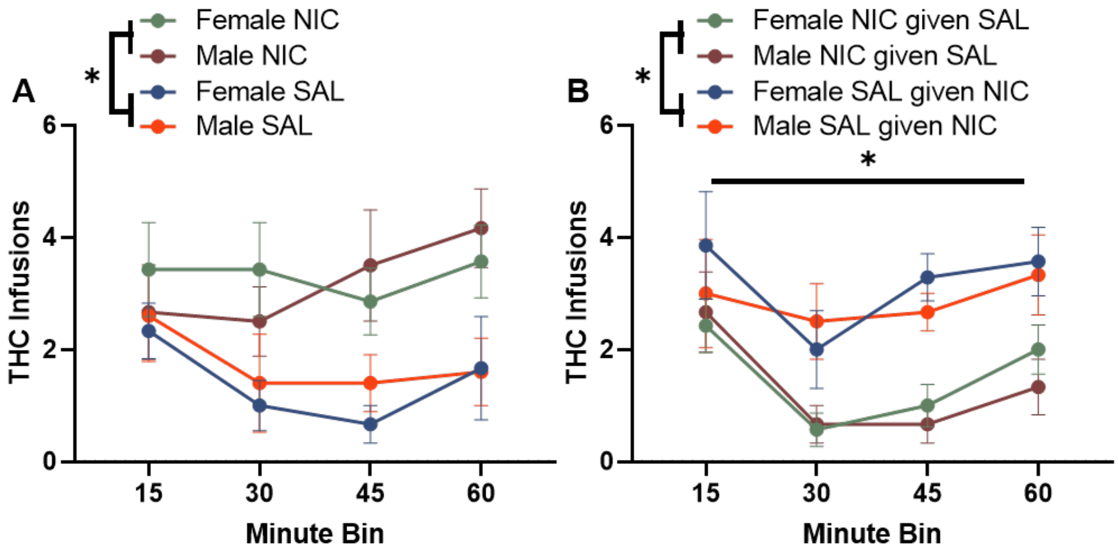


**Supplemental Figure S6**

Within session analysis of male and female rats self-administering THC during a 1-hour session divided into 15 minute bins. A) THC infusions received during each 15-minute bin on the last day of THC self-administration (day 22). A 3-way repeated measures ANOVA comparing infusions, drug injection, and sex resulted in a main effect of injection condition ($F_{(1, 20)} = 10.71$, $p<0.01$) but no other significant main effects or interactions ($p>0.1$ for all analyses). B) THC infusions received during each 15-minute bin on the last day of each animal receiving the opposite injection (day 17). There was a main effect of time ($F_{(3, 66)} = 6.95$, $p<0.01$) and a main effect of injection condition ($F_{(1, 22)} = 17.85$, $p<0.01$) but no other main effects or interactions emerged.
